# Urine proteome changes in a TNBS-induced colitis rat model

**DOI:** 10.1101/327080

**Authors:** Weiwei Qin, Ting Wang, Lujun Li, He Huang, Youhe Gao

**Affiliations:** Department of Biochemistry and Molecular Biology, Gene Engineering and Drug Biotechnology Beijing Key Laboratory, Beijing Normal University, Beijing 100875 China

## Abstract

Urine is an important resource for biomarker research. Without homeostasis, urine accumulates markers of all the changes in the body. Urine proteins reflect not only renal diseases but also changes in other organs in the body. However, urine has rarely been used to reflect inflammatory bowel disease. In the present study, a trinitrobenzene sulfonic acid (TNBS)-induced colitis rat model was used to mimic the human inflammatory bowel disease Crohn’s disease (CD). Urine samples from a control group (n=3), a TNBS 2-day group (n=3) and a TNBS 7-day group (n=3) were analyzed for candidate biomarker discovery by label-free and TMT-labeled proteomic quantitative methods. Seventy-seven urinary proteins were significantly changed in the colitis rats compared with that in the controls. These proteins were further validated by parallel reaction monitoring (PRM) targeted proteomic quantitative methods. Urine samples from the control group (n=8), the TNBS 2-day group (n=11) and the TNBS 7-day group (n=11) were analyzed by PRM. This led to the identification of 9 significantly differential expressed urinary proteins: CAH1, G3P, MMP-8, MANBA, NGAL, RNS1G, SLC31, S6A18, and TMM27. Based on the human protein tissue atlas, CAH1, RNS1G and SLC31 are highly enriched in the gastrointestinal tract. Among the 9 PRM-validated proteins, CAH1, MMP-8 and NGAL were previously reported as IBD-associated proteins (all exhibiting consistent trends with our observation), whereas the others are newly discovered by this study. Our results provide valuable clues for future study of urine biomarker of inflammatory bowel disease and Crohn’s disease.

## Introduction

Inflammatory bowel disease (IBD) is a group of chronic, relapsing intestinal inflammatory disorders with unknown etiology. The subtypes of IBD consist of Crohn’s disease (CD), ulcerative colitis (UC) and IBD unclassified (IBDU) (R). According to epidemiologic data, the incidence and prevalence of IBD are increasing with time in both high-income countries and newly industrialized countries [1, 2], indicating that it has become a global disease. In fact, 5% to 15% of cases are tentatively diagnosed as IBDU since no clear assignment for CD or UC can be made, and approximately 80% of those cases can later develop into UC or CD [3–5]. Current diagnosis and differential diagnosis mainly rely on clinical manifestations and endoscopic, radiological and histological criteria [6–8]. Endoscopy and imaging have notable limitations of cost, inconvenience, and invasiveness, making these procedures unsuitable for frequent monitoring of patients with IBD. Hence, there is a need for simple, noninvasive and accurate biomarkers for the diagnosis and prognosis of IBD diseases.

Mass spectrometry (MS)-based proteomics has dramatically improved and emerged as a prominent tool in the field of biomarker study. The development of isobaric tagging and targeted quantitative proteomic techniques makes the discovery of novel biomarkers feasible [9–12]. Many protein biomarkers of IBD have been described and are categorized as primarily blood and fecal biomarkers [13, 14]. Urine is an attractive resource for biomarker research that has been underutilized. Unlike blood, urine can accumulate markers of changes in the body without strict homeostatic regulation [15, 16]. Urinary proteomic studies have identified many candidate biomarkers for urogenital diseases, cardiovascular diseases and even cerebral diseases [17, 18]. However, there have been limited urinary biomarker studies for intestinal diseases, specifically for IBD, compared with the wide applications in renal diseases. Studying biomarkers in clinical samples remains challenging [19]. Studying the urine proteome is especially challenging because it is influenced by multiple factors, such as age, diet, exercise, gender, medication, and daily rhythms[20]. Animal models can be used to minimize the impact of many uncertain factors by establishing a direct relationship between a disease and corresponding changes in urine [21].

Candidate biomarkers of many diseases have been identified in urine using animal models. For example, significant changes in urine occurred in a unilateral ureteral obstruction rat model [22], and urinary proteins showed continuous changes with disease progression in an adriamycin-induced focal segmental glomerulosclerosis model [23]. The urinary proteome changed significantly before clinical symptoms and histopathological changes occurred in a bleomycin-induced pulmonary fibrosis model [24], an experimental autoimmune myocarditis model [25], a bacterial meningitis rat model [26], a glioblastoma animal model [27], and a Walker-256 tumor-bearing rat model [28].

Trinitrobenzene sulfonic acid (TNBS)-induced colitis is a hapten-induced colitis model that elicits a Th1-mediated immune response that involves IL-12 and TNF-α as cytokines [29, 30]. Histologically, transmural inflammation has been described with infiltrates of macrophages, neutrophils and lymphocytes, as well as colonic patch hypertrophy. From the immunological and histopathological characteristics, TNBS-induced colitis resembles features of CD [31].

This study was designed to identify a panel of urinary protein biomarkers related to CD using the TNBS-induced colitis rat model. Label-free and Tandem Mass Tags (TMT)-labeled quantitative proteomic methods were used for the discovery of biomarkers. Subsequently, candidates were validated by parallel reaction monitoring (PRM) targeted quantitative analysis using a quadrupole-orbitrap mass spectrometer.

## Materials and Methods

### Animals and Experimental Design

Male Wistar rats (180-200 g) were purchased from Charles River China (Beijing, China). All animals were maintained with a standard laboratory diet under controlled indoor temperature (21 ± 2 °C), humidity (65-70%) and 12 h light-dark cycle conditions. The animal experiments were reviewed and approved by the Peking Union Medical College (Approved ID: ACUC-A02-2014-008) and performed in accordance with the guidelines for animal research.

Experimental colitis was induced by TNBS (Sigma-Aldrich, St. Louis, MO, USA) according to the procedure described by Morris et al [32]. Briefly, rats were fasted for 24 h with water supplied. The next day (day 0), the rats were weighed and then anesthetized with ether. Under these conditions, 10 mg TNBS dissolved in 0.25 ml of 50% ethanol (n=18) or the same volume of 0.9% saline as a control (n=10) was administered rectally into the splenic flexure (6-8 cm from the anus) using a medical-grade polyurethane catheter (external diameter 2 mm). After instillation, a head-down position was maintained in order to prevent leakage and for even distribution of the hapten for 3 minutes. The bodyweight was measured daily. Animals were individually placed in rat fixators for 4 h to collect urine samples on day 2 and 7.

### Histopathology

For histopathology, the colon was harvested on day 2 (n=6) and day 7 (n=6) after rectal instillation. The colon was cut longitudinally, slightly cleaned in sterile water to remove fecal residues, and then fixed in 10% neutral-buffered formalin. The formalin-fixed tissues were embedded in paraffin, sectioned (4 mm) and stained with hematoxylin and eosin (HE) to reveal histopathological lesions.

### Urine sample preparation and tandem mass tag (TMT) labeling

Urinary proteins were extracted from the individual urine samples by acetone precipitation[33]. The urine samples were centrifuged at 3 000 g for 30 min at 4°C. After removing the cell debris, the supernatant was centrifuged at 12 000 g for 30 min at 4°C. Six volumes of acetone were added after removing the pellets, and the samples were precipitated at 4°C for 12 h. After centrifugation, the precipitates were dissolved in lysis buffer (7 M urea, 2 M thiourea, 120 mM dithiothreitol, and 40 mM Tris) and quantified by the Bradford method.

The proteins were digested with trypsin (Promega, USA) using filter-aided sample preparation methods [34]. Briefly, 200 μg of protein were loaded on a 10-kDa filter unit (Pall, USA). For digestion, the protein solution was reduced with 4.5 mM DTT for 1 h at 37°C and then alkylated with 10 mM indoleacetic acid for 30 min at room temperature in the dark. Finally, the proteins were digested with trypsin (1:50) for 14 h at 37°C. The resulting peptides were desalted and dried by a SpeedVac (Thermo Fisher Scientific, Waltham, MA, USA).

The nine urinary samples from the control group and the TNB-induced colitis group (Day 2, Day 7) were individually labeled with 126, 127, 127N, 128, 128N, 129, 129N, 130 and 131 TMT reagents according to the manufacturer’s protocol (Thermo Fisher Scientific, Germany) and then analyzed with two dimensional LC-MS/MS analysis.

### Off-line high-pH HPLC separation

The labeled peptide mixture was fractionated using off-line high-pH HPLC columns (XBridge, C18, 3.5 μm, 4.6 mm × 250 mm, Part No. 186003943; Waters, USA). The samples were loaded onto the column in buffer A (1 mM NH4OH in H2O, pH=10). The elution gradient was 5-30% buffer B (2 mM NH4OH in 90% acetonitrile, pH=10; flow rate=0.7 ml/min) for 60 min. The eluted peptides were collected at a rate of one fraction per minute. After lyophilization, the 60 fractions were resuspended in 0.1% formic acid and concatenated into 20 fractions (Zhao.MD).

### Online-LC-MS/MS analysis

Each peptide sample was dissolved in 0.1% formic acid and loaded on a trap column (75 μm × 2 cm, 3 μm, C18, 100 Å). The eluted gradient was 5-30% buffer B (0.1% formic acid in 80% acetonitrile; flow rate 0.3 μl/min) for 60 min. The eluent was transferred to a reversed-phase analytical column (50 μm × 150 mm, 2 μm, C18, 100 Å) by Thermo EASY-nLC 1200 HPLC system. Peptides were analyzed using Orbitrap Fusion Lumos mass spectrometer (Thermo Fisher Scientific, Germany). The MS data were acquired in the data-dependent acquisition mode. Survey MS scans were acquired in the Orbitrap using 350-1550 m/z range with the resolution set to 120 000. The most intense ions per survey scan (top speed mode) were selected for collision-induced dissociation fragmentation, and the resulting fragments were analyzed in the Orbitrap with the resolution set to 60,000 for labeled peptides and 30,000 for unlabeled peptides. Dynamic exclusion was employed with a 30 s window to prevent the repetitive selection of the same peptide. The normalized collision energy for HCD-MS2 experiments was set to 40% for labeled peptides and 30% for unlabeled peptides. Two technical replicate analyses were performed for each sample.

### Label-free quantitative LC-MS/MS data analysis

The raw MS data files were processed using Progenesis software (version 4.1, Nonlinear, Newcastle upon Tyne, UK) for label-free quantification[35], as previously described. Briefly, features with only one charge or more than five charges were excluded from the analyses. For further quantitation, all peptides (with Mascot score >30 and P<0.01) of an identified protein were included. Proteins identified by at least one peptide were retained. The MS/MS spectra were exported and processed with Mascot software (version 2.5.1, Matrix Science, London, UK) against the SwissProt rat database (released in July 2016, containing 7973 sequences). The following search parameters were used for protein identification: 10 ppm precursor mass tolerance, 0.02 Da fragment mass tolerance, up to two missed cleavage sites were allowed in the trypsin digestion, cysteine carbamidomethylation as affixed modification, and oxidation (M) as variable modifications. Only high confident peptide identifications with an FDR ⩽ 0.01 were imported into Progenesis LC-MS software for further analysis.

The statistical criteria of an ANOVAP value <0.05, a minimum of two peptides matched to a protein and a fold change >2 were used as the criteria for identification of differentially expressed proteins.

### TMT-abeled quantitative LC-MS/MS data analysis

The raw MS data files were searched using Proteome Discoverer (version 2.1; Thermo Fisher Scientific, San Jose, CA, USA) with Sequest HT against the SwissProt rat database (released in July 2016, containing 7973 sequences). The analysis workflow used included five nodes, namely, Spectrum Files (data input), Spectrum Selector (spectrum and feature retrieval), Sequest HT (sequence database search), Percolator (peptide spectral match or PSM Validation and FDR analysis), and Reporter Ions Quantifier (quantification). Sequest HT Search parameters were parent ion mass tolerance, 10 ppm; fragment ion mass tolerance, 0.05 Da; fixed modifications, carbamidomethylated cysteine (+58.00 Da) and labeled (+229.163 Da) N-terminal and lysine; and variable modifications, oxidized methionine (+15.995 Da). Other settings included default parameters. Unique and razor peptides were used for quantification values. Normalization mode is total peptide amount. All identified proteins had an FDR of ≤1%, which was calculated at the peptide level.

To be considered as being differentially expressed, proteins were required to have a p value <0.05 and a fold change >1.2.

### Parallel reaction monitoring (PRM) analysis

In the discovery phase, candidate protein biomarkers were identified using label-free and TMT-labeled quantitative proteomic methods. All the differentially abundant proteins were evaluated through targeted quantitation by PRM.

Pooled peptide samples (2 μg of each sample) were subjected to LC-MS/MS analysis (6 runs) for the spectrum library preparation. Skyline (Version 3.6.1 10279) [36]was used to build the spectrum library and filter peptides for PRM analysis. For each targeted protein, 2-6 associated peptides were selected using the following rules: (i) identified in the untargeted analysis with q value <1%, (ii) completely digested by trypsin, (iii) contained 8-18 amino acid residues, (iv) the first 25 amino acids at the N-terminus of proteins were excluded, and (v) fixed carbamidomethylation of cysteine. In total, 65 targeted proteins with 320 peptides were scheduled, and the RT segment was set to 4 min for each targeted peptide with its expected RT in the center based on the pooled sample analysis. The normalized collision energy was fixed to 30%, and a quadrupole isolation window of 0.7 Da was used to fragment the selected precursor ions. Prior to individual sample analysis, pooled peptide samples were subjected to PRM experiments to refine the target list. Finally, sixty-three proteins with 267 peptides could be targeted, as listed in Table S2.

Urine samples from the control group (n=8), the TNBS 2-day group (n=11), and the TNBS 7-day group (n=11) were used for the PRM analysis. All of the MS data were processed with Skyline. By comparing the same peptide across runs, the RT location and integration boundaries were adjusted manually to exclude interfering regions. Each protein’s intensity was quantitated using the summation of intensities from its corresponding transitions. Prior to the statistical analysis, the quantitated protein intensities were normalized by the summed intensity. The differential proteins were selected using one-way ANOVA, and p-values were adjusted by Benjamini & Hochberg method [37]. Significance was accepted at a p-value of less than 0.05.

## Result and discussion

### Clinical features of TNBS-induced Colitis in rats

Twenty-eight male Wistar rats (180-200 g) were divided into a control group (n=10) and a TNBS-treated group (n=18). Rats from the TNBS-treated group were given 10mg TNBS dissolved in 0.25 ml of 50% ethanol after ether anesthesia, and rats from the control group were given the same volume of 0.9% saline. The rats in the control group had normal daily activities, shiny hair and normal stool shape. In contrast, the rats in the TNBS-treated group showed hypomotility and piloerection, while the feces texture presented as dilute and mucous-like and continued to increase in severity to pus and blood. The bodyweight of the TNBS-treated rats suffered growth retardation and was significantly lower than that in the control group (p < 0.05) (Figure 2).

**Figure 1.**
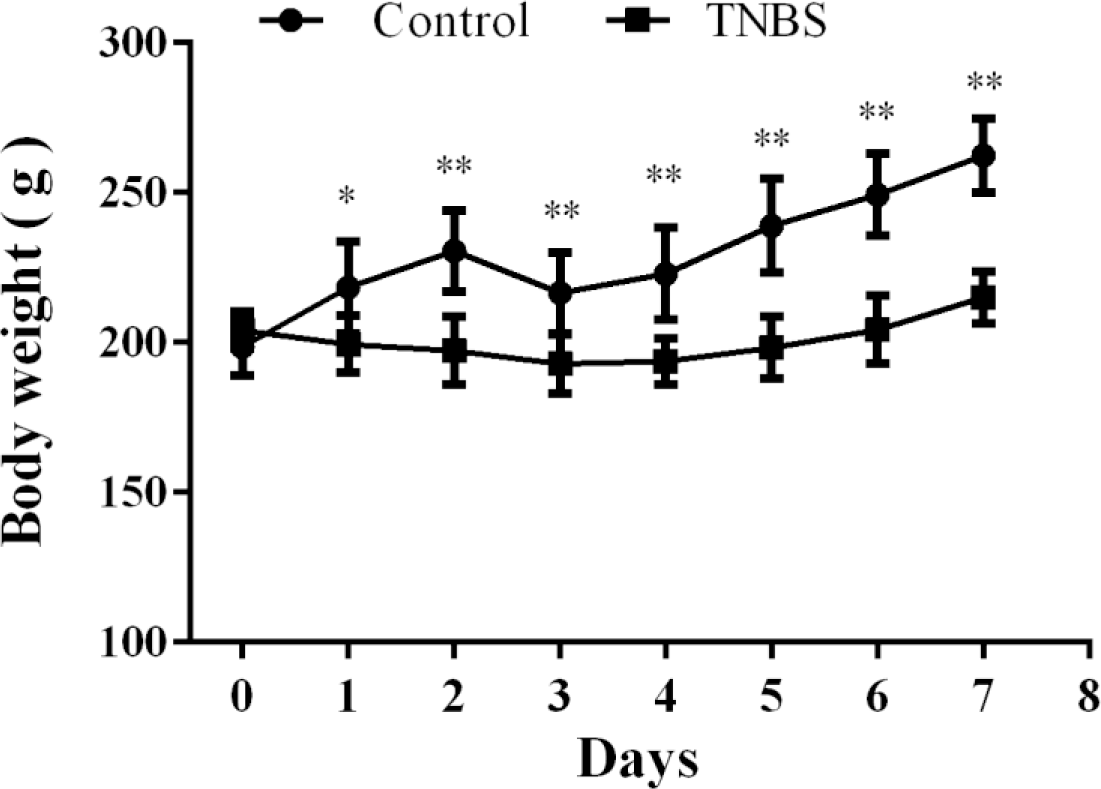
Bodyweight changes in the TNBS-induced colitis rat model. (n=11, * p<0.05, ** p<0.01)

**Figure 2.**
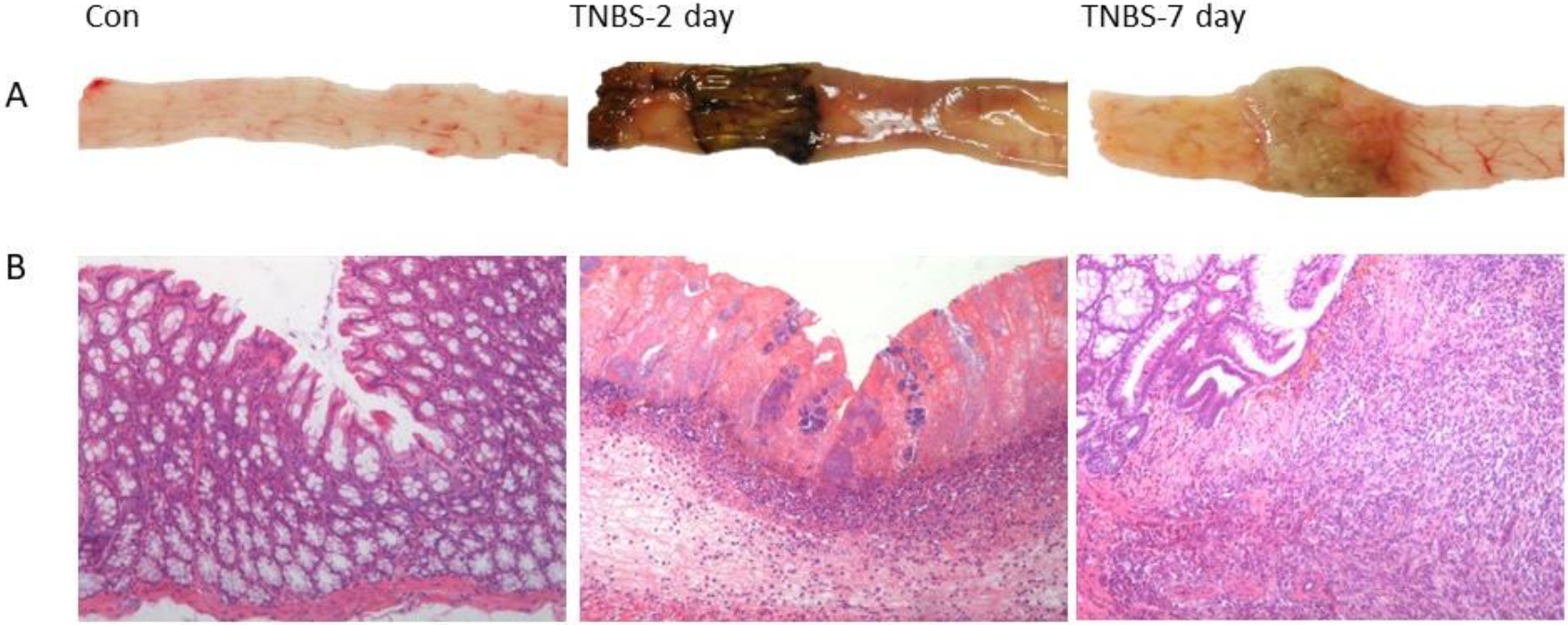
Histopathological characterization of TNBS-induced colitis. A:Macroscopic inspection, B: Hematoxylin and eosin stain (HE) at an original magnification 20X. (Con: Normal control rat colon; TNBS-2 day: TNBS treatment on day 2; TNBS-7 day: TNBS treatment on day 7)

The macroscopic and HE-stained microscopic features of the colon from control rats were typical and showed a normal structure: intestinal mucosa were clear and smooth (Fig. 3A); the epithelial cell layer was accompanied by the presence of goblet cells in straight tubular glands; and a normal quantity of cells were in the lamina propria (Fig. 3B). For the TNBS 2-day group, macroscopic inspection showed severe mucosal damage and hemorrhage (Fig. 3B). Under the microscope, the lesions could be seen and were mainly confined to the mucosa and submucosa with neutrophil infiltration and no tissue hyperplasia (Fig. 3B). For the TNBS 7-day group, macroscopic inspection showed prominent intestinal adhesion and colonic wall thickening (Fig. 3B), while inspection under the microscope showed diffuse inflammation extending through the muscularis mucosae, loss of the epithelium and goblet cells, and thickened lamina propria with the presence of fibroblasts.

**Figure 3.**
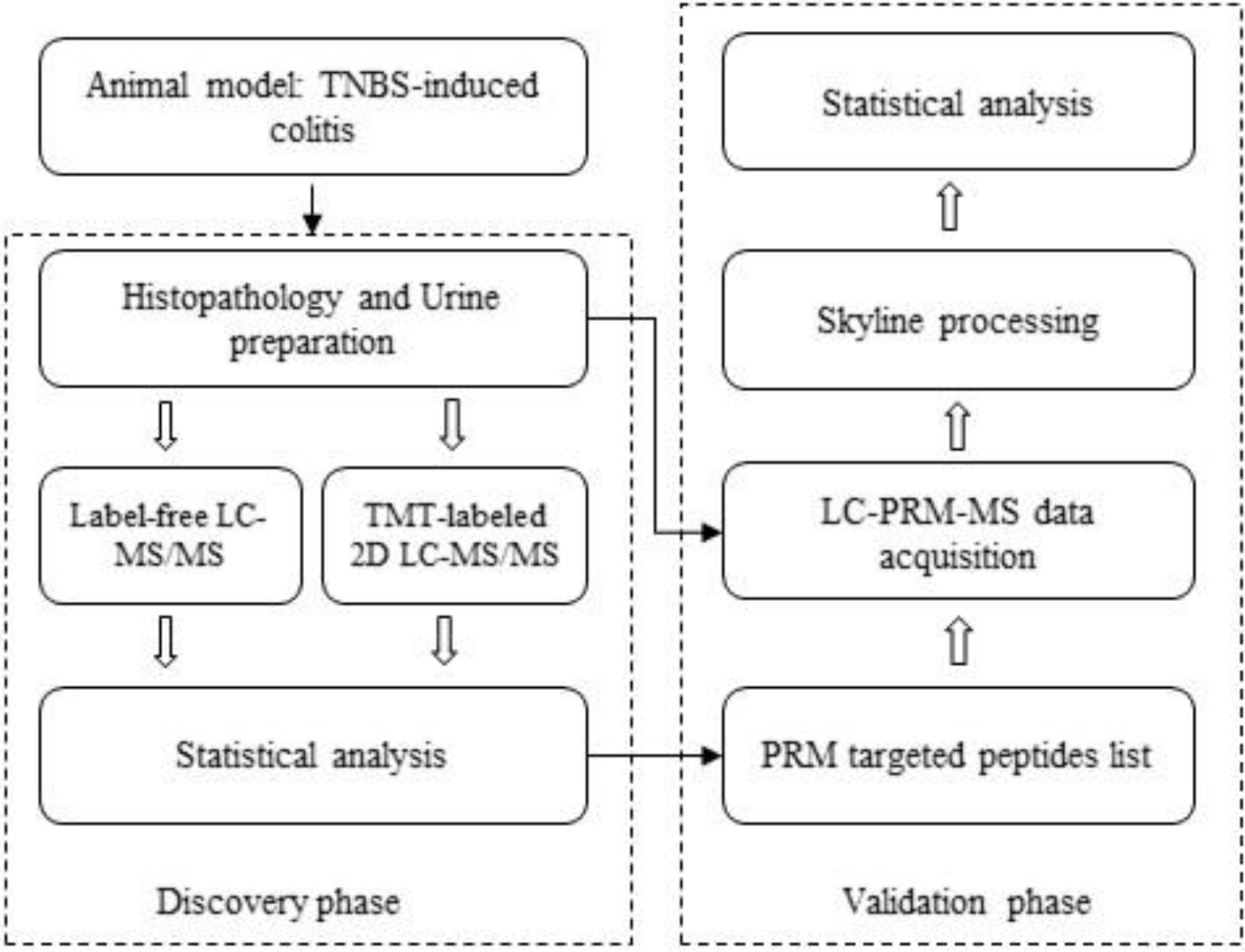
Workflow of the study of urine proteome changes in TNBS-induced colitis rats.

### Urine proteome changes

A summary of the overall experimental approach is presented in Figure 3. In the discovery phase, label-free and TMT-labeled proteomic quantitative methods were used to screen for changes in urinary proteins in TNBS-induced colitis rats. Urine samples from the control group (n=3), the TNBS-2 day group (n=3) and the TNBS-7 day group (n=3) were analyzed using these two methods.

In the label-free analysis, nine urine samples were analyzed by LC-MS/MS, with two technical replicates. The quantification was based on feature intensity using Progenesis software, and the database search performed by Mascot software. In total, 574 proteins with ≥2 unique peptides were identified with 1% FDR at the protein level. All identification and quantitation details are listed in Table S1. Compared to the control group, 22 and 32 proteins with significantly differential abundance (2-fold change, p<0.05) were identified in the TNBS-2 day and TNBS-7 day groups, respectively (Tables S2, S3).

In the TMT-labeled analysis, the same nine urine samples were labeled for offline two-dimensional reverse-phase liquid chromatography and high-resolution mass spectrometry (2D LC-MS/MS) analysis. The mixture was separated into 20 fractions by off-line high-pH HPLC, and then, each fraction was analyzed by LC-MS/MS two times. All spectra were processed by Proteome Discoverer (Version 2.1), and the quantification was based on the MS2 reporter. A database search yielded 49,417 high confidence peptides mapping to 1025 proteins with ≥2 unique peptides. All identification and quantitation details are listed in Table S4. Compared to the control group, 16 and 27 urinary proteins with significantly differential abundance (1.2-fold change, p<0.05) were identified in the TNBS-2 day and TNBS-7 day groups, respectively (Tables S5, S6).

### Parallel reaction monitoring (PRM) validation

After the label-free and TMT-labeled quantitative analyses, seventy-seven proteins (human homologous proteins) changed in the urine of TNBS-induced colitis rats. For further validation of these differential proteins, a PRM targeted proteomic quantitative method was used to analyze other urine samples from the control group (n=8), the TNBS-2 day group (n=11) and the TNBS-7 day group (n=11). In total, sixty-five targeted proteins with 320 peptides were screened. Prior to individual sample analysis, pooled peptide samples were subjected to PRM experiments in order to refine the target list. After further optimization, sixty-three proteins with 267 peptides were finally used for validation by PRM targeted proteomics (Table S7). The details of the targeted peptides are listed in Table S4. The technical reproducibility of the PRM assay was assessed, and the results show that among the targeted peptides, 229 peptides have abundance CV values that are less than 20% (Figure S1).

Overall, fifty-three of the 63 targeted proteins (25 increased and 28 decreased) exhibited the same average trend for the differential abundance of the proteins observed by both targeted and untargeted proteomic methods. Eighteen proteins show changes that were statistically significant (2-fold change, adjust p<0.05) (Table S8) on TNBS-2 day, and 9 proteins (4 increased and 5 decreased) show changes that were statistically significant on TNBS-7 day (Table 1) (Figure 4). Among the proteins showing significant changes, five proteins changed at both time points, and the trends were consistent. From the histopathology investigations, the second day of TNBS-induced colitis displayed mainly acute intestinal mucosal injury due to the effect of ethanol in the enema. The histopathological characteristics of the TNBS-7 day group resemble features of human immune inflammatory bowel disease (Crohn’s disease) [24]. Therefore, nine proteins showing differential abundance in the TNBS-7 day group have the potential to be urinary biomarkers of inflammatory bowel disease.

**Table 1.** Details of the changed urinary proteins identified by PRM on TNBS-7 day.

| UniProt ID | Protein Name | Day 2 |  | Day 7 |  | Human Ortholog |
| --- | --- | --- | --- | --- | --- | --- |
|  |  | FC | Adjust p | FC | Adjust p |  |
| Q8VD89 | Ribonuclease pancreatic gamma type (RNS1G) | 9.4 | 0.0013 | 12.4 | 0.0004 | P07998 |
| P30152 | Neutrophil gelatinase associated lipocalin (NGAL) | 29.8 | 0.0003 | 4.3 | 0.0429 | P80188 |
| O88766 | Matrix metalloproteinase-8 (MMP-8) | 2.7 | 0.0080 | 2.8 | 0.0052 | P22894 |
| P04797 | Glyceraldehyde-3-phosphate dehydrogenase (G3P) | 0.5 | 0.0216 | 0.5 | 0.0288 | P04406 |
| Q64319 | Solute carrier family 3 member 1 (SLC31) | 0.5 | 0.0362 | 0.4 | 0.0026 | Q07837 |
| Q62687 | Solute carrier family 6 member 18 (S6A18) | 0.6 | 0.1231 | 0.3 | 0.0004 | Q96N87 |
| Q4FZV0 | Beta-mannosidase (MANBA) | 1.4 | 0.0763 | 1.8 | 0.0328 | O00462 |
| Q9ESG3 | Collectrin (TMM27) | 0.7 | 0.4509 | 0.4 | 0.0262 | Q9HBJ8 |
| B0BNN3 | Carbonic anhydrase 1 (CAH1) | 1.0 | 0.9723 | 0.2 | 0.0026 | P00915 |

**Figure 4.**
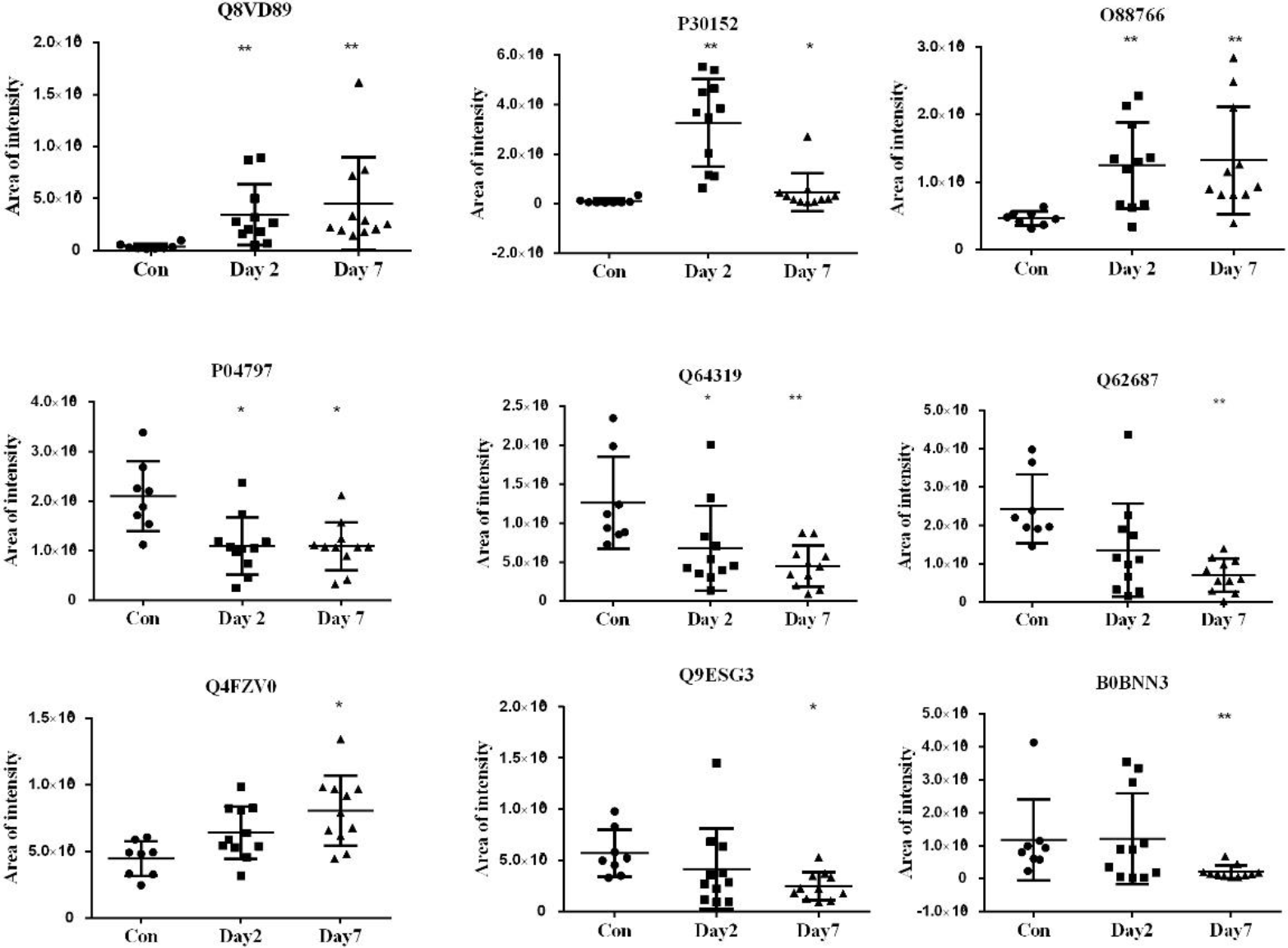
The intensity of significantly changed urinary proteins validated by PRM on TNBS-7 day. (ʪp<0.05, **p<0.01)

Among the 9 PRM-validated proteins in the TNBS-7 day group, three were highly enriched in the gastrointestinal tract based on the human protein tissue atlas [38]. These three are proteins are ribonuclease pancreatic gamma type (RNS1G), solute carrier family 3 member 1 (SLC31) and carbonic anhydrase 1(CAH1). Additionally, three the TNBS 7-day group PRM-validated proteins had been reported as biomarkers of inflammatory bowel disease. Neutrophil gelatinase-associated lipocalin (NGAL) has been shown to be significantly increased in the urine of Crohn’s disease patients compared to baseline samples[39]. It is also found to be upregulated in the serum and fecal samples of CD patients [40]. Matrix metalloproteinase-8 (MMP-8) is reported to be significantly elevated in plasma in patients with inflammatory bowel disease compared to that in normal controls[41]. Carbonic anhydrase 1(CAH1) is reported to be observed at lower levels in the urine of patients with Crohn’s disease and ulcerative colitis compared to that found in a normal control group [42].

There were also some differential proteins discovered in our study that have never been reported to be related to inflammatory bowel disease, such as glyceraldehyde-3-phosphate dehydrogenase (G3P), solute carrier family 6 member 18 (S6A18), beta-mannosidase (MANBA), and collectrin (TMM27), as well as RNS1G and SLC31. Since these proteins were dramatically changed, they also have the potential to be urinary biomarkers of inflammatory bowel disease.

In this preliminary study, only a TNBS-induced colitis animal model was used to discover urinary biomarkers. Further analysis of other animal models may provide more sensitive and specific candidate biomarkers that may be capable of discriminating between Crohn’s disease (CD) and ulcerative colitis (UC). In addition, a large number of clinical samples should be used to verify specific proteins or protein patterns as clinically applicable biomarkers of inflammatory bowel disease.

## Conclusion

In this study, nine candidate urinary biomarkers of IBD were uncovered in a TNBS-induced colitis rat model. Three of these biomarkers had been previously reported to be related to IBD. Additionally, three were annotated as highly enriched in the gastrointestinal tract. Our study showed that urine can be a good source of IBD biomarkers. To further illustrate the application of our identifications, urine samples from IBD patients will be included in further clinical studies. To validate these nine urinary proteins in clinical samples, targeted proteomic approaches will be used in the future.

## Acknowledgment

National Key Research and Development Program of China (2016YFC1306300); Beijing Natural Science Foundation (7173264, 7172076); the Fundamental Research Funds for the Central Universities (2015KJJCB21); Beijing cooperative construction project (110651103); Beijing Normal University (11100704).

